# Insights into the Interactions of Peptides with Monolayer-Protected Metal Nanoclusters

**DOI:** 10.1101/2022.11.27.518090

**Authors:** Vikas Tiwari, Sonali Garg, Tarak Karmakar

**Affiliations:** Department of Chemistry, Indian Institute of Technology, Delhi, Hauz Khas 110016, New Delhi, India

## Abstract

Monolayer-protected atomically precise metal nanoclusters (MPC) are an important class of molecules that have potential applications in catalysis, imaging, and drug delivery. Recent studies have shown that peptide-based drugs can be complexed with MPCs to avoid enzymatic degradation and get delivered to targeted cells. Although the MPCs potential role in imaging and drug delivery processes have been studied, for their impactful use, specific molecular interactions between MPCs and biomolecules, mainly proteins and peptides should be explored in detail. In this work, we have carried out atomistic molecular dynamics simulations to investigate the interactions between Au-based MPCs and an anticancer peptide, melittin. The MEL peptides get attached to the MPCs surface by the formation of multiple hydrogen bonds between the peptide amino acid residues with MPCs ligands. Additionally, the positively charged residues such as Lys and Arg, the Trp, and the N-terminal of the peptide anchor strongly to the MPC core playing a crucial role in the peptide’s overall stabilization on the MPC surface.

---

Monolayer protected metal clusters, abbreviated as MPCs, are a class of molecules having a metal (mostly noble metals) core and an organic ligand layer that aids in stabilizing the metal nanostructure.^1-2^ Unlike metal nanoparticles with sizes ranging from 1 to 100 nm and lacking atomic accuracy (do not have well-defined molecular formula), MPCs are atomically accurate (*‘precise’*) nanoclusters with a tunable size range of 1-3 nm,^3-5^ which means they have a fixed ratio of metal to the ligand. Due to their size dependent properties and changeable ligand layers, they are exploited in the design of nanostructured materials. Exceptional catalytic activity, multivalency,^6-8^ high luminescence, electrical and optical properties, good photostability,^9^ cooperative binding, and biocompatibility are just a few of their many desirable characteristics. The ligand functionalization of metal clusters in the MPCs makes them applicable for biological sensing, multi-modal imaging of cancer cells, and therapeutic activity.^10^ Inside the cell, MPCs get stabilized by interacting with various biomolecules which makes them suitable for targeted cell imaging. Au-nanoclusters (Au-NCs) have a self-illuminating property used in optical fluorescence imaging as they give a contrasting graph.^11-12^ Functionalized Au-NCs act as therapeutic agents such as GSH-protected Au-NCs as radio-sensitizers for cancer radiotherapy and TAT-peptide Au-NCs for photodynamic therapy.^10^ MPCs can also be used as drug carriers for various biomedical agents such as photosensitizers, and chemotherapeutics. The MPCs stabilize the drug molecules on their surface by specific intermolecular interactions and effectively deliver them to targeted sites thereby preventing severe side effects due to non-specific delivery.^13^ Till now, a variety of MPCs with different metal cores such as Au, Ag, and Pd using thiolate ligands have been synthesized and characterized.^14-16^ Among them, the thiolate-protected gold nanoclusters having molecular formula Au_25_(SR)_18_ are the most studied ones, both theoretically and computationally, due to their early discovery, easy synthesis, and potential applications in the field of nano-bio-medicine.^17-20^

In MPCs, the covalent Au-S bonding interactions stabilize the metal core, and the ligands help the MPCs to interact with other molecules such as protein residues, nucleic acids, lipids, and other biological fluids.^21-24^ Proteins interact through corona formation with the MPCs and show various non-covalent interactions such as van der Waals, hydrophobic, and hydrogen bonding interactions.^25-27^ Because of these interactions, proteins undergo several conformational changes causing unfolding, leading to proteins aggregates, and proteins-Au-NC assemblies. Qi et al. have studied the complexation of peptides with MPCs^28^ and opened up the possibility of using MPCs as drug carriers. Lately, peptide-based drugs have emerged to be an important class of therapeutics that are promising for the treatment of various types of tumors. However, one concern is that many of such peptides get degraded and proteolyzed in the biological environment.^29^ Use of MPCs is beneficial in such a case as MPCs provide extra stabilization to the peptides by anchoring them on their surface. Additionally, they help facile penetration to target tissues and the controlled delivery of the drugs. The binding of proteins/peptides to the Au-NCs is governed by Au-NCs size, the hydrophobicity and the hydrophilicity of the functionalized ligands. These peptide-MPC interactions are crucial for anti-cancer activities by lyses, releasing infected ions and molecules in the cell cytoplasm, eventually leading to cell apoptosis.

The combination of biomolecules and nanoclusters has been used for many applications in the past years.^30-34^ A study by Arora *et al*. showed that a recombinant protein PTEN bound to silver nanoclusters and coated with polymer PEG, resulted in the reduction of cell viability of MCF7 cells.^35^ In another study, Xia *et al*. found that fluorescence imaging and chemo-photodynamic therapy near-infrared region for tumor-targeted cells using polypeptide-modified gold nanoclusters (GNCs)-based nanoprobes, inhibits the growth of tumor cells.^36^ Significant efforts have been made to understand the mechanism of binding biomolecules with metal nanoclusters.^37-41^ Despite all these works, a proper understanding of the mechanism and the kind of interactions responsible for the formation of MPC-bio complexes is still limited. In fact, Antoine *et al*., in their recent comment, discussed the need for computational works in the study of protein-NC interactions.^42^

In this work, we have investigated the interactions between the 26 amino acid residues long peptide melittin (MEL) and the MPC Au_25_(MHA)_18_ in an aqueous medium using atomistic molecular dynamic (MD) simulations. A series of equilibrium MD simulations of systems with various combinations of MPC and MEL have been carried out to characterize the type of interactions and rationalize the enhanced stabilization of MEL on the MPC surface.

To study the interactions responsible for the formation of a complex between the melittin (MEL) and Au_25_(MHA)_18_ (Au-MPC), systems with different ratios of MEL and Au-MPC were prepared using the configurations obtained from individual simulations (see Table 1). The idea behind taking different ratios of MEL:Au-MPC was to mimic systems with dilute and concentrated solutions of the MEL and Au-MPC mixtures. We simulated two sets of systems. The first set has the following compositions - (i) MEL : Au25_MPC (1:1), (ii) MEL:Au-MPC (1:2), (iii) MEL:Au-MPC (2:1), and (iv) MEL(dimer):Au-MPC (2:1). In another set, we simulated (i) a system containing 10 MEL and 10 Au-MPC and (ii) another one with the ratio 10:20:10 of MEL-monomer, MEL-dimer, and Au-MPC, respectively. Details of the systems, simulation parameters, and computational methods are discussed in the supporting information (SI).

## MEL:Au-MPC (1:1) complex

In the 1:1 MEL:Au-MPC system, both MEL and MPC obtained from individual simulations (see section X of the supporting information (SI)) were kept close to each other in a cubic box filled with water. Within a few tens of nanoseconds, the two molecular entities come together and form a MEL:AuMPC complex which remains intact throughout the entire simulation duration. The driving force for the complex formation between MEL and Au-MPC is the formation of hydrogen bonds between the carboxylic acid groups of the MHA ligand and the polar MEL residues (Arg, Lys, and GLN) present at the C-terminal. Subsequently, the Au-MPC’s ligands start interacting with the peptide backbone. Once the MEL is properly attached to the surface, its Lys21 sidechain penetrates through the ligands and interacts with the metal surface. The positively charged -NH_3_ group either interacts with the metal core or the partially negatively charged sulfur atoms of the ligands. This residue has been found to get anchored to the metal core for the simulation duration of 500 ns (Fig. 2a) The probability distribution of the distance between Lys21’s nitrogen and the sulfur atom of one of the ligands with which it shows a dominant peak at distance 0.34 nm and a smaller one at 0.42 nm. This indicates that the Lys21 sidechain shuttles between two sites - one near the chosen S atom and the other one near an Au atom (see Fig. 2b)

**Figure 1:**
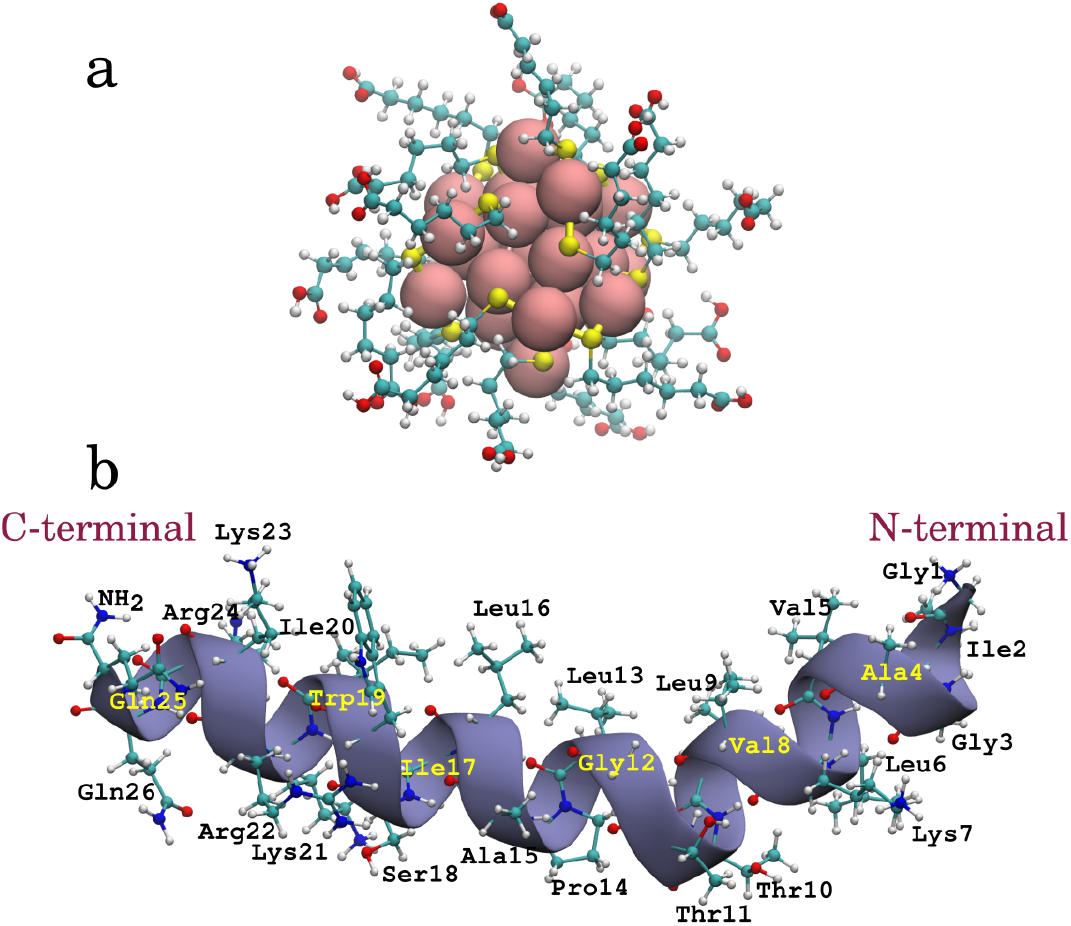
(a) 3D structure of an Au-MPC - Au_25_(MHA)_18_ and (b) 3D structure of melittin protein monomer with labeled amino acid residues.

**Figure 2:**
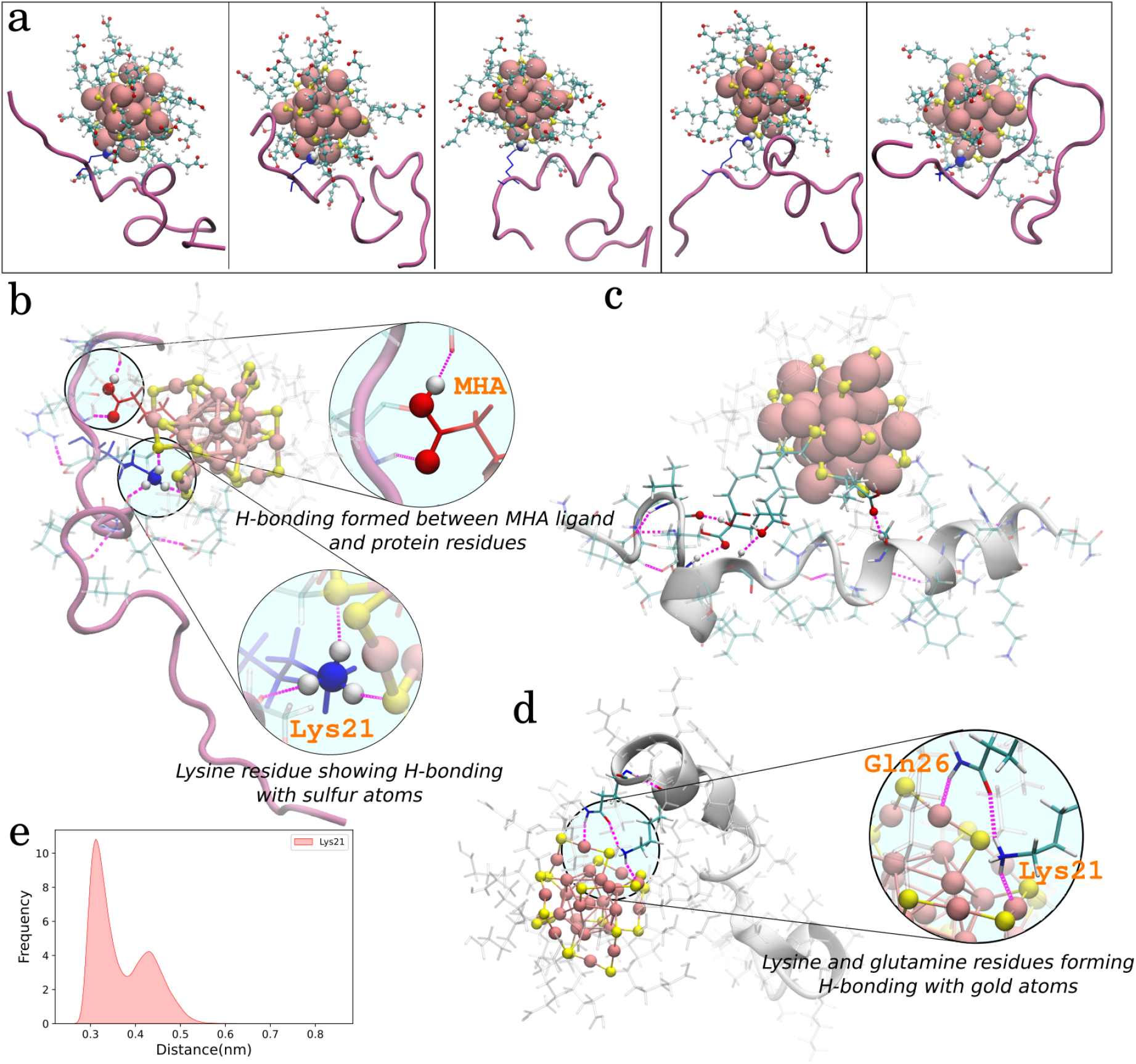
1MEL:1MPC simulation analysis, a) MEL showing various conformations after getting anchored via its Lys21 residue, b) Hydrogen bonding interactions showing by Lys21 with sulfur atoms, c) initial interactions showing various hydrogen bonds between MHA ligand and protein residues, d) Lys21 and Gln26 showing a network of hydrogen bonding with surface gold atoms and e) Lys21 residues distance distribution with respect to one of the sulfur atoms.

We now switch our attention to the conformational dynamics of MEL anchored on the MPC. The root mean square deviations (RMSD) of MEL backbone atoms, root mean square fluctuations of residue heavy atoms, hydrogen bonding analysis, and residue-residue distance distributions were calculated to quantify the conformational flexibility of the anchored peptide and delineate specific intermolecular interactions.. The RMSD profile shown in Fig. 3a indicates that initially the peptide has large conformational flexibility, and once it gets anchored to the MPC surface the fluctuations have been suppressed. This manifests enhanced stabilization of the peptide on the MPC surface. Furthermore, from the calculated RMSF for all the residues shown in Fig. 3b, it is evident that the region where the Lys21 is present shows the least fluctuations. Additionally, Ile17, Ser18, and Ile20 residues also show comparatively lower fluctuations than other residues. This may be due to the hydrophobic interactions between these residues with the ligands’ aliphatic chains. The other Lys7 residue near the N-terminal is exposed to the solution and thus has a relatively larger RMSF value than the Lys21. Apart from Lys21, Gln26 also stabilizes the MEL: AuMPC complex by interacting with the metal core (Fig. 2d). Formation of hydrogen bonding is also one of the major interactions observed in this system, hence we calculated the hydrogen bonds between the MPC ligands and the MEL protein (Figure 3c). On average 3-5 hydrogen bonds are formed providing extra stability to the MEL-MPC complex.

**Figure 3:**
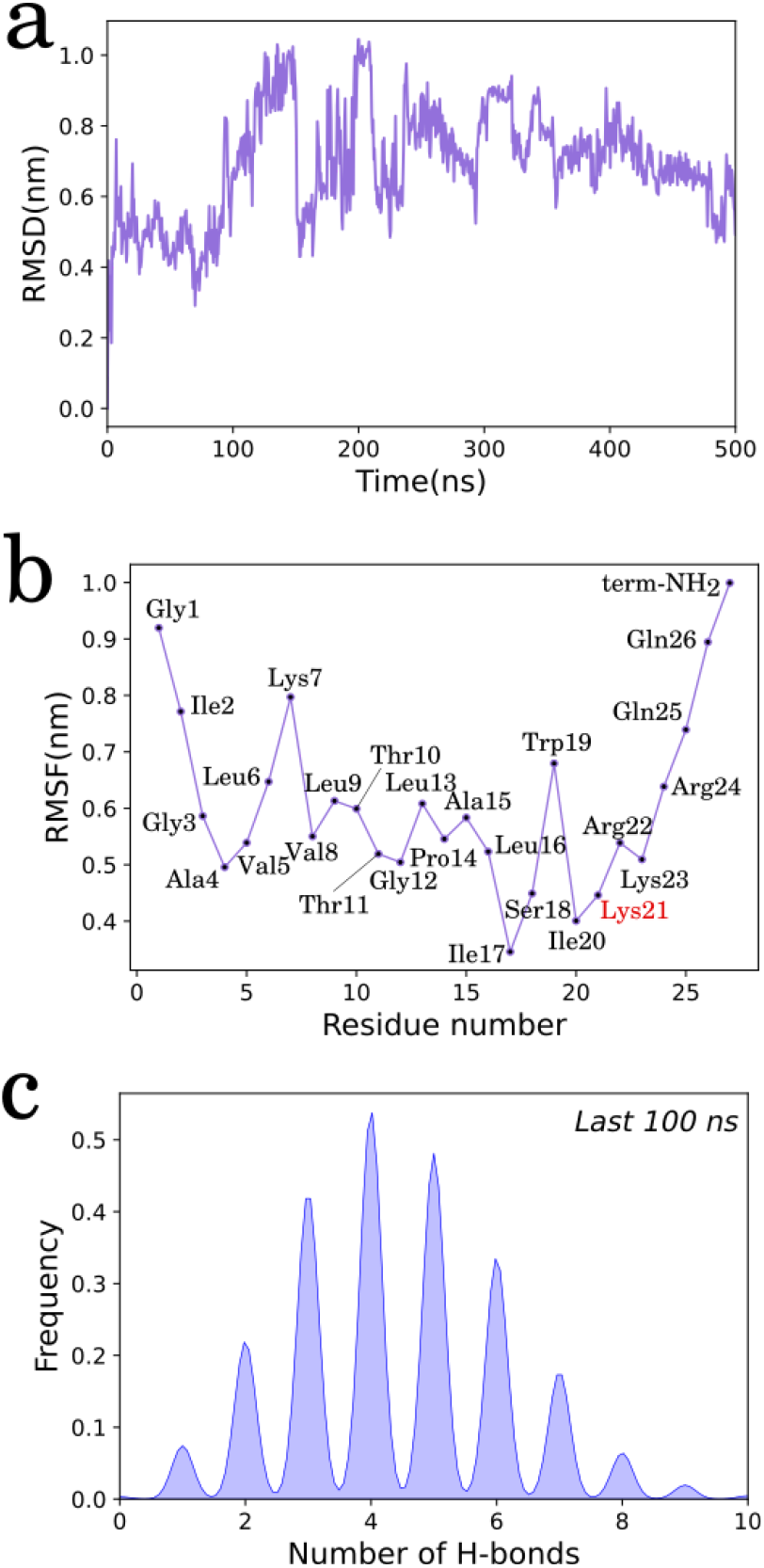
1MEL:1MPC simulation analysis, a) RMSD, b) RMSF, and c) hydrogen bonding analysis between MEL and MHA ligands

## MEL:Au-MPC (1:2) complex

Understanding the 1:1 system of MEL:MPC has given an interesting perception of the MEL-MPC interactions but left us with some important questions to focus upon. We saw that only one Lys was able to get inside the MPC, why not two or more? One of the reasons might be the geometrical feasibility of the interaction. More importantly, the stability of the MEL on the MPC surface in the 1:1 system is not quite good as it shows fluctuations throughout the simulation. So we anticipated, if there was another MPC in the system, this may lead to better stability of the MEL protein. Hence, to test our hypothesis, we simulated a 1:2 MEL-MPC system keeping two MPCs and a MEL in close proximity (Fig. 4). In particular, we have placed one additional MPC to the MEL:AuMPC 1:1 complex. It is not surprising that the interaction between Lys21 and the metal core remains intact in this system as well. Within a few nanoseconds of the simulation, the second MPC (MPC2) has been found to interact with the MEL through the formation of hydrogen bonds between peptide residues and the ligands carboxylic acid groups. The MEL’s orientation makes it possible to interact with the MPC2, and in this orientation, its positively charged N-terminal can access the MPC2 metal surface. We have gained some fascinating understandings from this serendipitous orientational attachment of MEL with a pair of MPCs. The ligands in Au-MPCs are orientated in a certain manner, causing areas of low ligand density, which prevents the Au-MPCs from being spherical. The possibility of protein residues becoming attached to these sites is increased by the existence of these patches on MCPs, which provide an accessible path to the metal surface. The trajectory visualization demonstrated that primarily the metal surface near the Gly1 had high ligand density which prevented the interactions between them. After a while, the MPC rotated in such a way that the low ligand density side was exposed near the Gly1 residue. After that Gly1 residue entered the exposed area of the metal surface and started to interact via hydrogen bonding. Figure 4 shows images illustrating the mechanism through which Gly1 residue is anchored to the surface of gold.

**Figure 4:**
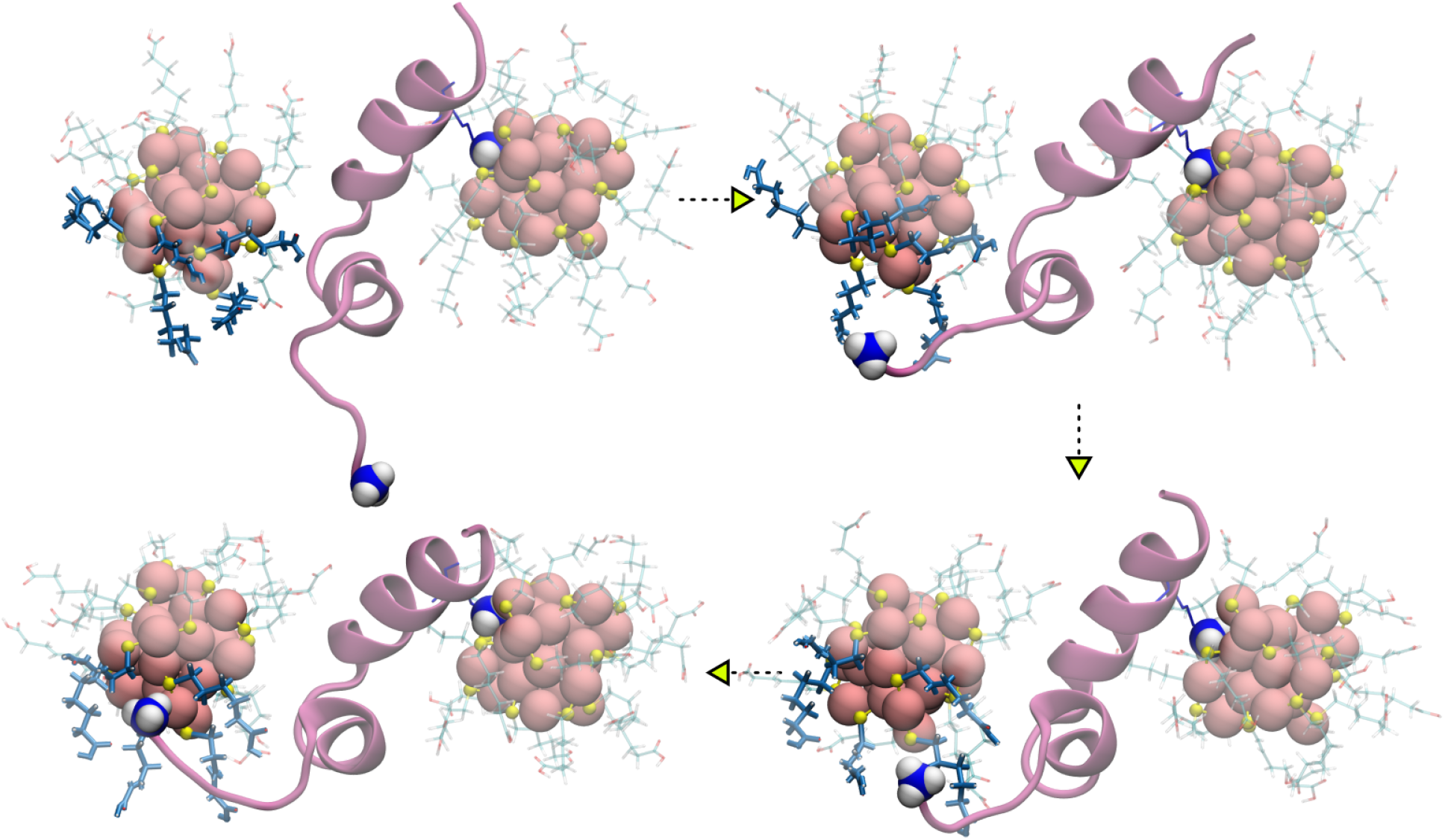
Mechanism of N-terminal interaction with 2nd MPC (MPC2) molecule in 1:2 MEL:Au-MPC simulation.

The anchoring of two MPCs at two terminals of the MEL led to an ‘S’-shaped locked conformation which remained stable for 500 ns simulation time. In this conformation, the peptide shows suppressed conformational dynamics compared to that in the case of the 1:1 system (Fig. 5a). To further quantify its stability, RMSF analysis was performed for the last 100 ns simulation trajectory (Fig. 5e). This analysis showed that most of the residues of MEL have very little fluctuations that are less than 0.25 nm. This gives a clear indication of improved stability of MEL in presence of two MPCs. Along with Lys21 and Gly1, few other residues were seen interacting with the metal surface for some time (Fig. 5c). Furthermore, to identify the residues that come in contact with the MPC surface, a distance analysis was performed for all the residues (Fig. 5d). The distance distributions show a large peak of Lys21 and Gly1 near 0.35 nm distance indicating the anchoring of these residues with the MPCs. Whereas, Arg24 and Trp19 show smaller peaks below 0.5 nm distance suggesting that these residues only interact for a short time interval. Interestingly, Trp19 shows a large peak near 1.4 nm which was later found due to the hydrophobic interaction of Try19 aromatic rings with MHA’s hydrophobic region. Additionally, a closer look at the simulation trajectory revealed that when Lys21 interacts with the metal surface, the non-polar residues such as Leu13, Pro14, Ala15, Leu16, and Ile17 form hydrophobic interactions with the MPCs ligands providing extra stability to the conformation. As observed in the 1:1 system, here also the hydrogen bonding interactions, on an average of 5-6 hydrogen bonds (Fig. 5b), between the peptide residues and MPC ligands carboxylic acid groups are retained which provide additional stabilization.

**Figure 5:**
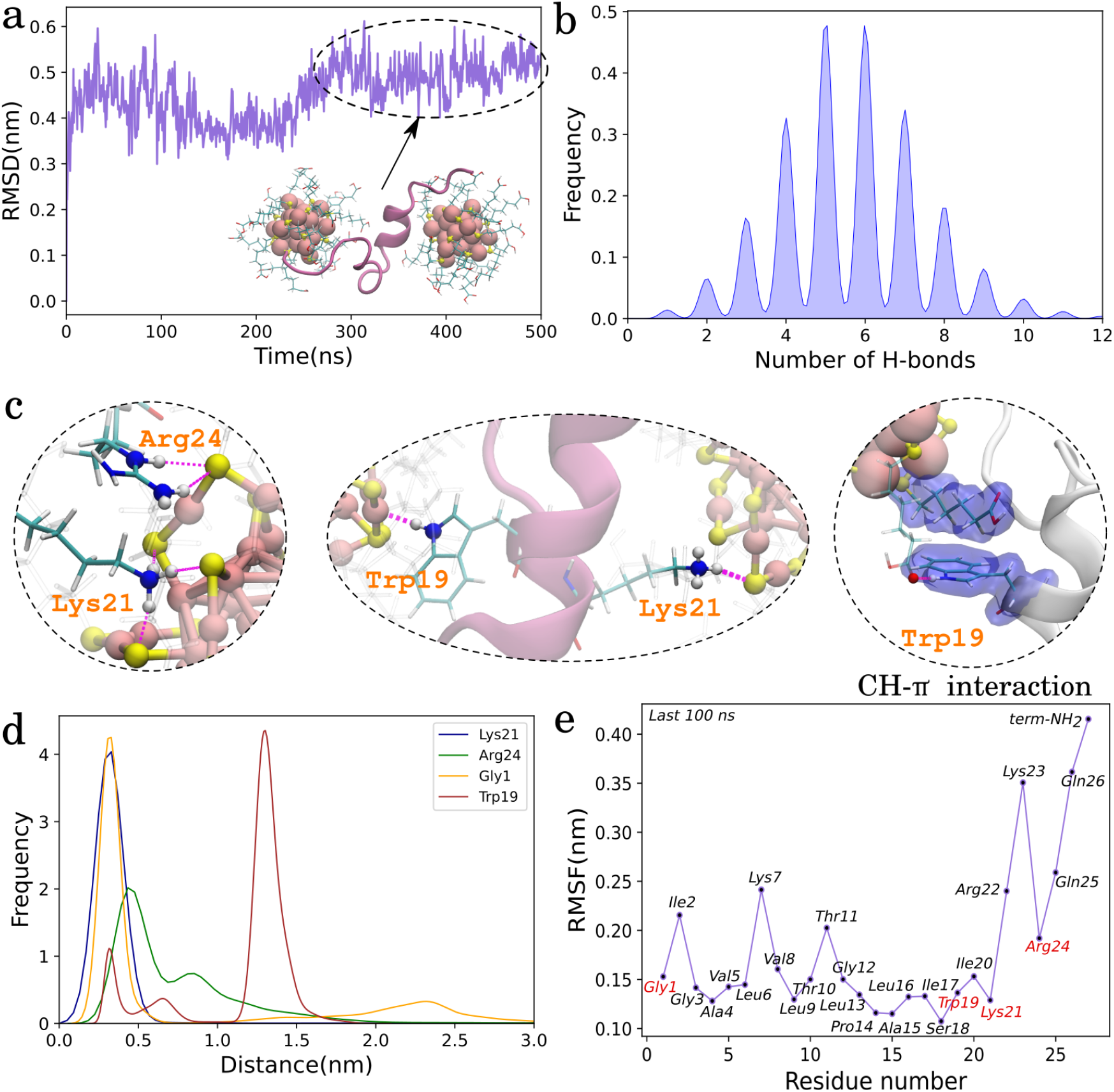
1MEL-2MPC simulation analysis, (a) RMSD, (b) hydrogen bond distribution between MEL residues and MHA ligands, (c) Various interactions observed during the simulation, (d) distance distribution analysis of various residues interacting with metal surface, and (e) RMSF analysis of last 100 ns trajectory.

## MEL:Au-MPC(2:1) complex

We saw in the 1:2 system the presence of one extra MPC exerts additional stabilization of the MEL on the MPCs surface. Here we explore another possibility in which we place two MEL peptides in close proximity to a single MPC molecule. Our aim is twofold: we are interested in finding if (i) one MPC can carry two MEL peptides, and (ii) whether the binding of one MEL on a MPC influences the other one or not. At the start, the two MEL monomers were far apart from the MPC. Within 10 ns simulation time, the Lys23 (near the C-terminal) of MEL2, and the Thr11 (near the N-terminal) of MEL1, began to interact *via* hydrogen bonding with the MPCs metal surface along with a few other residues forming hydrogen bonding with MPC ligands. These interactions were only present for a short time mainly responsible for bringing the peptides and the MPC close together. Subsequently, at the latter simulation time, a stable conformation was obtained. In this structure, we observed the formation of three non-conventional hydrogen bonding interactions - (i) MEL1-Arg24, (ii) MEL1-Gly3 (backbone NH), and (iii) MEL2-Trp19 which also showed CH-*π* interactions with MPC ligands (Fig. Sx). Additionally, many intermolecular hydrogen bonds were present between MEL1 and MEL2 residues and MPC ligands. These hydrogen bonds stabilize the peptides on the MPC surface. RMSD analysis of both the MEL1 and MEL2 protein chain shows very low fluctuations for the last 100 ns (Fig. 6c) indicating suppressed conformational dynamics of the two peptides (Fig. 6b). To check on the fluctuations of MEL residues in the *‘locked conformational’* state, RMSF analysis was performed for the last 100 ns of the simulation. From Fig. 6d, it can be found that most of the residues have RMSF values <0.25 nm, and only MEL1’s C-terminal residues Gln25, Gln26, and NH_2_ (C-terminal) show fluctuations up to 0.45 nm (Fig. 6c). Further distance analysis was performed for the residues who were found to be interacting with the metal surface. The histogram plot of the distance analysis showed that MEL1-Gly3, MEL2-Lys23 and MEL2-Trp19 showed a peak near 0.3-0.4 nm indicating a good interaction with the metal surface via hydrogen bonding (Fig ref ESI). Whereas MEL1-Arg24 showed a broad peak at 0.25-0.75 nm due to the presence of 2 NH2 and 1 NH group which can have a variety of arrangements during the hydrogen bonding formation. MEL1-Gly3, MEL2-Lys23, and MEL2-Trp19 also showed a peak after 0.5 nm which indicates long interaction of these residues with the MPC ligands at some different locations as observed from the visualization of the trajectory.

**Figure 6:**
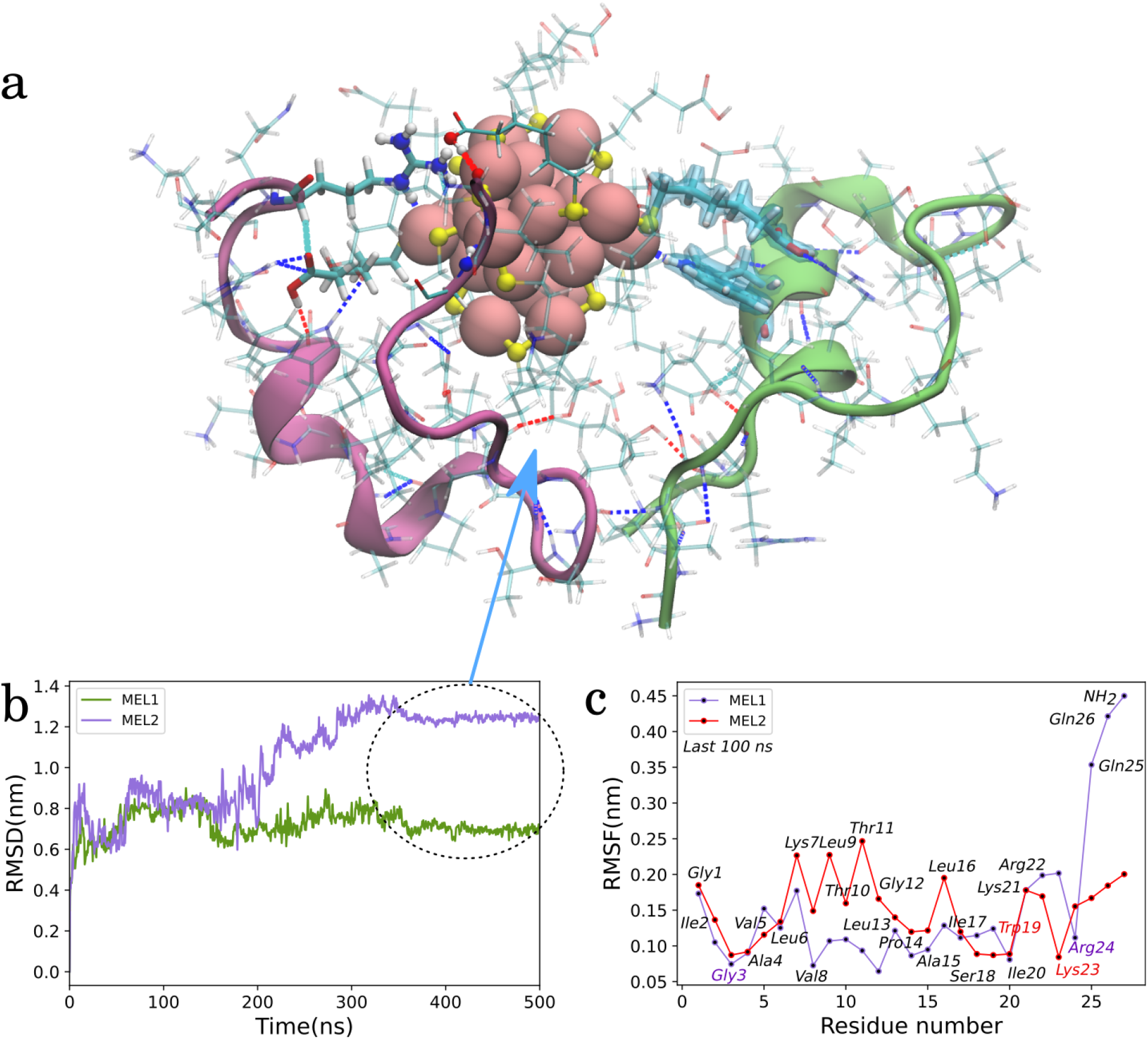
2MEL:1MPC simulation analysis, a) a stable conformation of 2MEL-1MPC complex obtained in the last 100 ns of simulation showing various interactions, b) RMSD analysis of both MEL, c) RMSF analysis of last 100ns trajectory.

MEL has been found to exist in many multimeric forms depending upon various conditions such as concentration, ionic strength, temperature, and pH. So the 2MEL-1MPC system can be also formed if a dimer-MEL interacts with the MPC. Hence to have a complete understanding we first simulated a MEL dimer in water for 100 ns. A configuration collected from this simulation was placed to an MPC, and the system was simulated for 500 ns. One of the interesting observations from this simulation was the interaction of the MEL with MPC via a hydrophobic patching (Fig 7). As the dimeric MPC approached the MPC, an exposed surface on the MPC was developed and the dimer-MEL got attached to it. We conjecture that this interaction may be relevant to MEL’s anticancer activity where the exposed positively charged residues play an important role in breaking the membrane of cancerous cells. Further analyses for this system have been discussed in ESI section xx.

**Figure 7:**
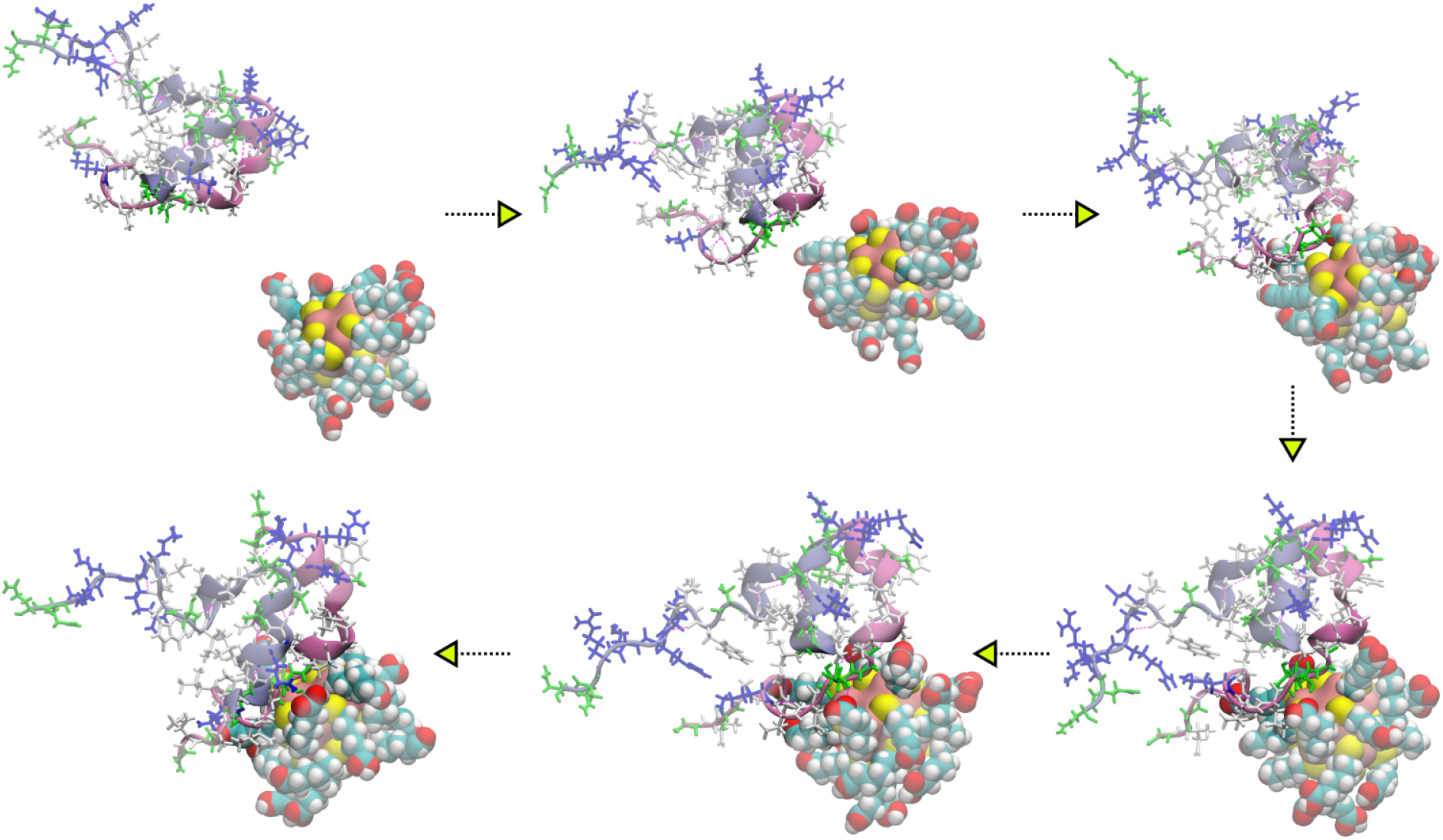
Mechanism showing adsorption of dimeric melittin on the exposed gold surface of MPC in the dimer-MEL:MPC (1:1) simulation.

**Figure 8:**
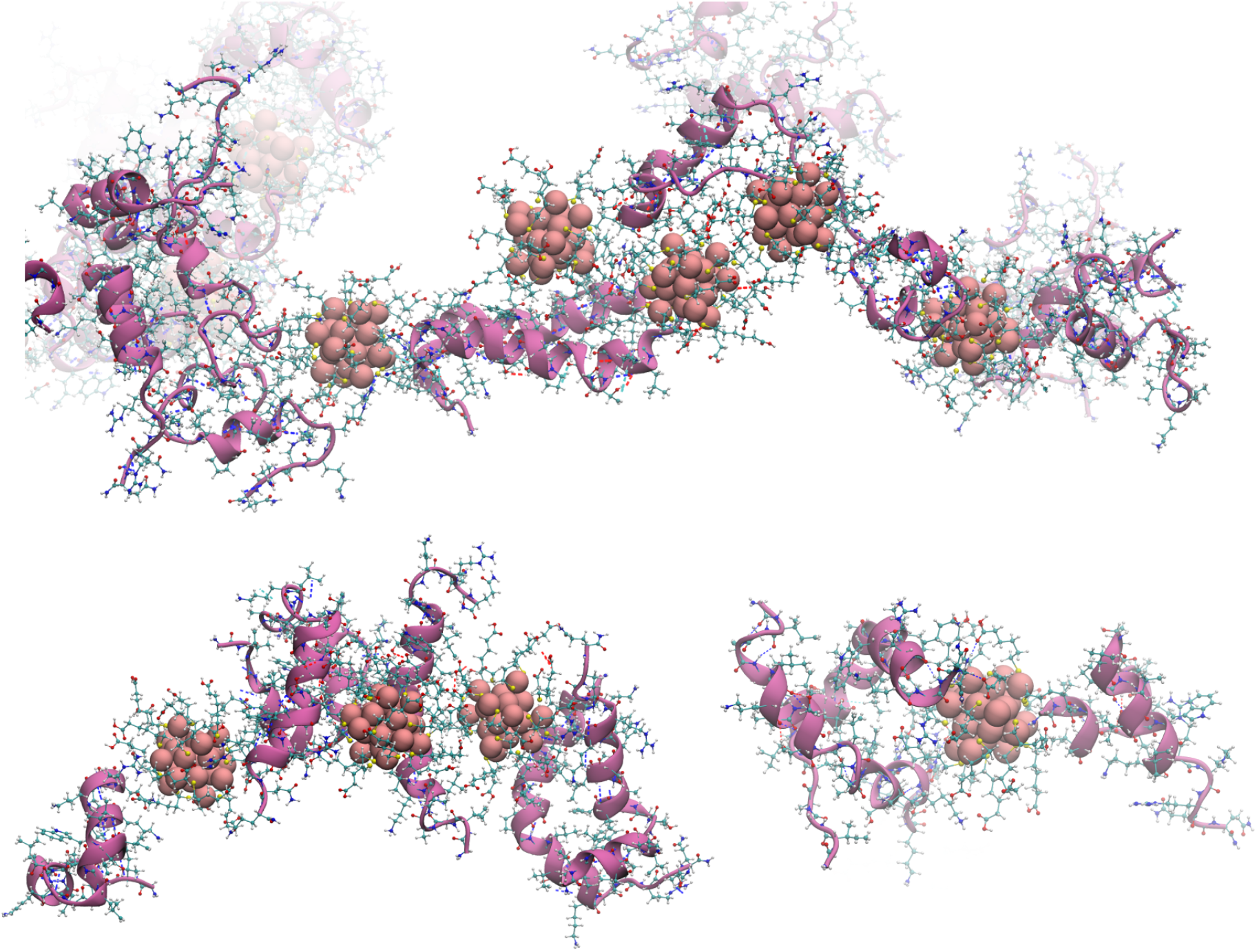
Interactions of monomeric and dimeric MEL with multiple MPCs

## MEL:MPC (10:10) system

The systems we have discussed so far represent dilute solutions of MPC and MEL. To understand their interaction patterns in concentrated solution, we simulated two more systems - (i) in one, we have randomly placed 10 MEL monomers and 10 MPC, and (ii) in another, 10 monomeric and 5 dimeric MEL and 10 MPC kept in a large cubic box. The purpose of this simulation was to understand the behavior of these molecules present in a denser environment where there are possibilities of forming bigger clusters. As expected, in both simulations, we have observed the formation of MPC clusters as well as MEL aggregation. Both dimeric and monomeric MEL peptides were found to get attached to the MPC cluster.

In summary, the results obtained from various simulations of MEL-MPC complexes have provided a better understanding of interactions responsible for the MEL-MPC complex formation. The stabilization effect has been found to be due to a combination of many favorable interactions such as (i) the hydrogen bonding interactions between the peptides’ polar residues with the carboxylic acid group of the ligands on the MPC surface, (ii) strong interactions (electrostatic and hydrogen bonding) between positively charged sidechains (Lys and Arg) as well as the N-terminal with the metal core and the sulfur atoms of the ligands on the metal surface, and (v) interactions between non-polar residues with the ligands hydrophobic patches on the MPC. One of the important interactions which was observed during the simulation was the non-conventional hydrogen bonding between polar residues of MEL and the surface gold atoms. This type of interaction has been reported previously using ab initio DFT calculations between the polar amino acid residues and Au_3_ system.^43^ This strong interaction of Au-MPC with the MEL may be the reason for its stability in the biological environment and the other free positively charged residues might be responsible for carrying out the lysis of cell membranes of the cancerous cells maintaining its anti-cancer activity.

## COMPUTATIONAL DETAILS

The initial structure of a mercaptohexanoic acid (MHA) ligand was generated using Avogadro software, and the complete MPC structure and topology were generated using the NanoModeler server^44^ which utilizes general AMBER force field (GAFF)^45^ for parametrizing MPCs. The starting structure of melittin was obtained from the RCSB PDB data bank. Amber99sb-ildn force field^46^ was used to generate the topology of the melittin. The system was solvated using the TIP3P water molecules, and it was neutralized using chloride ions. All the simulations were performed using GROMACS version 2021.4.^47^

For calculating the long-range electrostatic interactions, the Particle Mesh Ewald (PME) with grid spacing 0.16 and order 4 was used. After the systems had been completely neutralized, the system’s energy was minimized by employing the steepest descent minimization technique. Each system was brought to an equilibrium temperature of 300 K using the NVT ensemble. The system’s pressure at 1 bar was maintained by equilibrating the system using an NPT ensemble using the Berendsen barostat.^48^ Both NVT and NPT equilibration simulations were performed for 250 ps using a time step of 0.5 fs without any constraint. The NVT and NPT MD simulations were carried out with the help of a Leap-Frog integrator. The V-rescale modified Berendsen thermostat was utilized to successfully couple the temperatures with a temperature coupling constant of 0.5 ps. For pressure coupling in the production NPT simulations, the isotropic Parrinello-Rahman algorithm^49^ was employed with a coupling constant of 2 ps. The value for the short-range van der Waals cutoff and the short-range electrostatic cutoff was set to 1.0 nm each. Finally, each system was for a total of 500 ns in the NPT ensemble, and the integration was performed using a time step of 2 fs. During the NPT production run, all h-bonds were constrained using the LINCS algorithm.^50^

The VMD software package^51^ was used for the visualization of simulation trajectories and the preparation of figures. The GROMACS tools were used to perform the RMSD, RMSF, and Hydrogen bond analyses, while the PLUMED version 2.8.0^52^ was used to perform the contact analysis of protein residues with the MPC. All the analyses graphs were generated using in-house Python codes.

## Notes

### Competing Interest Statement

The authors have declared no competing interest.

## Bibliography

1) Jin, R.; Zeng, C.; Zhou, M.; Chen, Y. Atomically Precise Colloidal Metal Nanoclusters and Nanoparticles: Fundamentals and Opportunities. Chemical Reviews 2016, 116 (18), 10346–10413.

2) Chakraborty, I.; Pradeep, T. Atomically Precise Clusters of Noble Metals: Emerging Link between Atoms and Nanoparticles. Chemical Reviews 2017, 117 (12), 8208–8271.

3) Malola, S.; Nieminen, P.; Pihlajamäki, A.; Hämäläinen, J.; Kärkkäinen, T.; Häkkinen, H. A Method for Structure Prediction of Metal-Ligand Interfaces of Hybrid Nanoparticles. Nature Communications 2019, 10 (1).

4) Nag, A.; Pradeep, T. Assembling Atomically Precise Noble Metal Nanoclusters Using Supramolecular Interactions. ACS Nanoscience Au 2022, 2 (3), 160–178.

5) Malola, S.; Häkkinen, H. Prospects and Challenges for Computer Simulations of Monolayer-Protected Metal Clusters. Nature Communications 2021, 12 (1).

6) Longo, A.; Boed, E. J. J.; Mammen, N.; Linden, M.; Honkala, K.; Häkkinen, H.; Jongh, P. E.; Donoeva, B. Towards Atomically Precise Supported Catalysts from Monolayer-Protected Clusters: The Critical Role of the Support. Chemistry – A European Journal 2020, 26 (31), 7051–7058.

7) Sudheeshkumar, V.; Sulaiman, K. O.; Scott, R. W. J. Activation of Atom-Precise Clusters for Catalysis. Nanoscale Advances 2020, 2 (1), 55–69.

8) Shi, Q.; Qin, Z.; Sharma, S.; Li, G. Recent Progress in Heterogeneous Catalysis by Atomically and Structurally Precise Metal Nanoclusters. The Chemical Record 2021, 21 (4), 879–892.

9) Tlahuice-Flores, A.; Whetten, R. L.; Jose-Yacaman, M. Ligand Effects on the Structure and the Electronic Optical Properties of Anionic Au_25_(SR)_18_ Clusters. The Journal of Physical Chemistry C 2013, 117 (40), 20867–20875.

10) Song, X.-R.; Goswami, N.; Yang, H.-H.; Xie, J. Functionalization of Metal Nanoclusters for Biomedical Applications. The Analyst 2016, 141 (11), 3126–3140.

11) Su, Y.; Xue, T.; Liu, Y.; Qi, J.; Jin, R.; Lin, Z. Luminescent Metal Nanoclusters for Biomedical Applications. Nano Research 2019, 12 (6), 1251–1265.

12) Qu, X.; Li, Y.; Li, L.; Wang, Y.; Liang, J.; Liang, J. Fluorescent Gold Nanoclusters: Synthesis and Recent Biological Application. Journal of Nanomaterials 2015, 2015, 1–23.

13) Rizvi, S.; Saleh, A. King Abdulaziz Medical City, National Guard Health Affairs, Mail Code 6610. King Abdullah International Medical Research Center (KAIMRC) 2142, 9515.

14) Zhang, B.; Sels, A.; Salassa, G.; Pollitt, S.; Truttmann, V.; Rameshan, C.; Llorca, J.; Olszewski, W.; Rupprechter, G.; Bürgi, T.; Barrabés, N. Ligand Migration from Cluster to Support: A Crucial Factor for Catalysis by Thiolate-Protected Gold Clusters. ChemCatChem 2018, 10 (23), 5372–5376.

15) Si, K. J.; Chen, Y.; Shi, Q.; Cheng, W. Nanoparticle Superlattices: The Roles of Soft Ligands. Advanced Science 2017, 5 (1), 1700179.

16) Li, Y.; Zhai, T.; Chen, J.; Shi, J.; Wang, L.; Shen, J.; Liu, X. Water-Dispersible Gold Nanoclusters: Synthesis Strategies, Optical Properties, and Biological Applications. Chemistry – A European Journal 2021, 28 (10).

17) Kumar, S.; Jin, R. Water-Soluble Au25(Capt)18 Nanoclusters: Synthesis, Thermal Stability, and Optical Properties. Nanoscale 2012, 4 (14), 4222.

18) Parker, J. F.; Fields-Zinna, C. A.; Murray, R. W. The Story of a Monodisperse Gold Nanoparticle: Au_25_L_18_. Accounts of Chemical Research 2010, 43 (9), 1289–1296.

19) Zhu, M.; Lanni, E.; Garg, N.; Bier, M. E.; Jin, R. Kinetically Controlled, High-Yield Synthesis of Au_25_ Clusters. Journal of the American Chemical Society 2008, 130 (4), 1138–1139.

20) Kang, X.; Chong, H.; Zhu, M. Au_25_(SR)_18_: The Captain of the Great Nanocluster Ship. Nanoscale 2018, 10 (23), 10758–10834.

21) Goswami, N.; Zheng, K.; Xie, J. Bio-NCs – the Marriage of Ultrasmall Metal Nanoclusters with Biomolecules. Nanoscale 2014, 6 (22), 13328–13347.

22) Bhattacharya, S. R.; Bhattacharya, K.; Xavier, V. J.; Ziarati, A.; Picard, D.; Bürgi, T. The Atomically Precise Gold/Captopril Nanocluster Au_25_(Capt)_18_ Gains Anticancer Activity by Inhibiting Mitochondrial Oxidative Phosphorylation. ACS Applied Materials & Interfaces 2022, 14 (26), 29521–29536.

23) Brancolini, G.; Corazza, A.; Vuano, M.; Fogolari, F.; Mimmi, M. C.; Bellotti, V.; Stoppini, M.; Corni, S.; Esposito, G. Probing the Influence of Citrate-Capped Gold Nanoparticles on an Amyloidogenic Protein. ACS Nano 2015, 9 (3), 2600–2613.

24) Matus, M. F.; Malola, S.; Häkkinen, H. Ligand Ratio Plays a Critical Role in the Design of Optimal Multifunctional Gold Nanoclusters for Targeted Gastric Cancer Therapy. ACS Nanoscience Au 2021, 1 (1), 47–60.

25) Tavanti, F.; Pedone, A.; Menziani, M. C.; Alexander-Katz, A. Computational Insights into the Binding of Monolayer-Capped Gold Nanoparticles onto Amyloid-β Fibrils. ACS Chemical Neuroscience 2020, 11 (19), 3153–3160.

26) Shemetov, A. A.; Nabiev, I.; Sukhanova, A. Molecular Interaction of Proteins and Peptides with Nanoparticles. ACS Nano 2012, 6 (6), 4585–4602.

27) Wang, P.; Wang, X.; Wang, L.; Hou, X.; Liu, W.; Chen, C. Interaction of Gold Nanoparticles with Proteins and Cells. Science and Technology of Advanced Materials 2015, 16 (3), 034610.

28) Qi, J.; Liu, Y.; Xu, H.; Xue, T.; Su, Y.; Lin, Z. Anti-Cancer Effect of Melittin-Au25(MHA)18 Complexes on Human Cervical Cancer HeLa Cells. Journal of Drug Delivery Science and Technology 2022, 68, 103078.

29) Lau, J. L.; Dunn, M. K. Therapeutic Peptides: Historical Perspectives, Current Development Trends, and Future Directions. Bioorganic & Medicinal Chemistry 2018, 26 (10), 2700–2707.

30) Yu, X.; Dai, Y.; Zhao, Y.; Qi, S.; Liu, L.; Lu, L.; Luo, Q.; Zhang, Z. Melittin-Lipid Nanoparticles Target to Lymph Nodes and Elicit a Systemic Anti-Tumor Immune Response. Nature Communications 2020, 11 (1).

31) Cantarutti, C.; Raimondi, S.; Brancolini, G.; Corazza, A.; Giorgetti, S.; Ballico, M.; Zanini, S.; Palmisano, G.; Bertoncin, P.; Marchese, L.; Patrizia Mangione, P.; Bellotti, V.; Corni, S.; Fogolari, F.; Esposito, G. Citrate-Stabilized Gold Nanoparticles Hinder Fibrillogenesis of a Pathological Variant of β_2_-Microglobulin. Nanoscale 2017, 9 (11), 3941–3951.

32) Jallouk, A. P.; Palekar, R. U.; Marsh, J. N.; Pan, H.; Pham, C. T. N.; Schlesinger, P. H.; Wickline, S. A. Delivery of a Protease-Activated Cytolytic Peptide Prodrug by Perfluorocarbon Nanoparticles. Bioconjugate Chemistry 2015, 26 (8), 1640–1650.

33) Muhamad, N.; Plengsuriyakarn, T.; Na-Bangchang, K. Application of Active Targeting Nanoparticle Delivery System for Chemotherapeutic Drugs and Traditional/Herbal Medicines in Cancer Therapy: A Systematic Review. International Journal of Nanomedicine 2018, Volume 13, 3921–3935.

34) Tang, M.; Gandhi, N. S.; Burrage, K.; Gu, Y. Interaction of Gold Nanosurfaces/Nanoparticles with Collagen-like Peptides. Physical Chemistry Chemical Physics 2019, 21 (7), 3701–3711.

35) Arora, N.; Gavya S L.; Ghosh, S. S. Multi-Facet Implications of PEGylated Lysozyme Stabilized-Silver Nanoclusters Loaded Recombinant PTEN Cargo in Cancer Theranostics. Biotechnology and Bioengineering 2018, 115 (5), 1116–1127.

36) Xia, F.; Hou, W.; Zhang, C.; Zhi, X.; Cheng, J.; de la Fuente, J. M.; Song, J.; Cui, D. PH-Responsive Gold Nanoclusters-Based Nanoprobes for Lung Cancer Targeted Near-Infrared Fluorescence Imaging and Chemo-Photodynamic Therapy. Acta Biomaterialia 2018, 68, 308–319.

37) Tavanti, F.; Pedone, A.; Menziani, M. C. Disclosing the Interaction of Gold Nanoparticles with Aβ(1–40) Monomers through Replica Exchange Molecular Dynamics Simulations. International Journal of Molecular Sciences 2020, 22 (1), 26.

38) Dutta, S.; Corni, S.; Brancolini, G. Molecular Dynamics Simulations of a Catalytic Multivalent Peptide–Nanoparticle Complex. International Journal of Molecular Sciences 2021, 22 (7), 3624.

39) Tavanti, F.; Pedone, A.; Menziani, M. C. Multiscale Molecular Dynamics Simulation of Multiple Protein Adsorption on Gold Nanoparticles. International Journal of Molecular Sciences 2019, 20 (14), 3539.

40) Charchar, P.; Christofferson, A. J.; Todorova, N.; Yarovsky, I. Understanding and Designing the Gold–Bio Interface: Insights from Simulations. Small 2016, 12 (18), 2395–2418.

41) Brancolini, G.; Bellucci, L.; Maschio, M. C.; Di Felice, R.; Corni, S. The Interaction of Peptides and Proteins with Nanostructures Surfaces: A Challenge for Nanoscience. Current Opinion in Colloid & Interface Science 2019, 41, 86–94.

42) Antoine, R.; Maysinger, D.; Sancey, L.; Bonačić-Koutecký, V. Open Questions on Proteins Interacting with Nanoclusters. Communications Chemistry 2022, 5 (1).

43) Pakiari, A. H.; Jamshidi, Z. Interaction of Amino Acids with Gold and Silver Clusters. The Journal of Physical Chemistry A 2007, 111 (20), 4391–4396.

44) Franco-Ulloa, S.; Riccardi, L.; Rimembrana, F.; Pini, M.; & De Vivo, M. NanoModeler : A Webserver for Molecular Simulations and Engineering of Nanoparticles. Journal of Chemical Theory and Computation 2018, 15 (3), 2022–2032.

45) Wang, J., Wolf, R. M.; Caldwell, J. W.; Kollman, P. A.; Case, D. A. “Development and testing of a general AMBER force field”. Journal of Computational Chemistry 2004, 25, 1157–1174.

46) Lindorff-Larsen, K.; Piana, S.; Palmo, K.; Maragakis, P.; Klepeis, J. L.; Dror, R. O.; Shaw, D. E. Improved side-chain torsion potentials for the Amber ff99SB protein force field. Proteins: Struct., Funct., Genet. 2010, 78, 1950– 1958.

47) Berendsen, H. J.; van der Spoel, D.; van Drunen, R. GROMACS: a message-passing parallel molecular dynamics implementation. Comput. Phys. Commun. 1995, 91, 43– 56.

48) Berendsen, H. J.; Postma, J. P. M.; Gunsteren, W. F.; DiNolam, A.; Haak, J. R. Molecular dynamics with coupling to an external bath. J. Chem. phys. 1984, 81, 3684.

49) Parrinello, M.; Rahman, A. “Polymorphic transitions in single crystals: A new molecular dynamics method,” J. Appl. Phys., 1981, 52, 7182–7190.

50) Hess, B.; Bekker, H.; Berendsen, H. J.; Fraaije, J. G. E. M. “LINCS: A linear constraint solver for molecular simulations,” J. Comp. Chem., 1997, 18, 1463–1472.

51) Humphrey, W.; Dalke, A.; Schulten, K. VMD - Visual Molecular Dynamics. J. Molec. Graphics, 1996, 14, 33–38.

52) Tribello, G. A.; Bonomi, M.; Branduardi, D.; Camilloni, C.; Bussi, G. PLUMED2: New feathers for an old bird. Comp. Phys. Comm. 2014, 185, 604.

