## Supplemental Information for "Insights into the Interactions of Peptides with Monolayer-Protected Metal Nanoclusters"

**Table 1: Systems simulated**

| | **Ratio of Mel to MPCs** | **System size (number of atoms)** |
|---|---|---|
| MPC = Au25(MHA)18 | | |
| **Monomeric Mel** | 1:1 | 33623 |
| | 1:2 | 98238 |
| | 2:1 | 98277 |
| | 10:10 | |
| **Dimeric Mel (See ESI)** | 2:1 | 98241 |
| **Both mono and dimeric Mel** | 30:10<br>10(mono) & 10(di) | |

### Mel in water

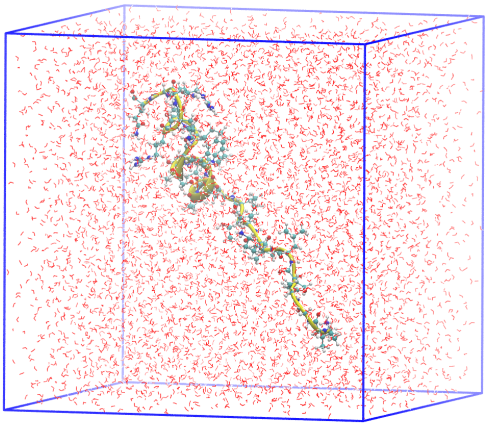

### MPC in water

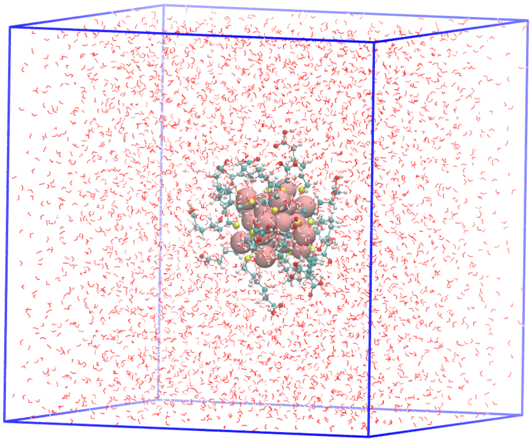

***Figure S1:*** *Individual MEL and MPC in water simulations*

**Dimer Mel:MPC(1:1)**

Melittin is also found to be stable in the dimeric form in certain conditions. So, we simulated dimeric melittin in water for 100ns. Then a 1:1 dimeric MEL and $Au_{25}$-MPC system was simulated. In this both MEL and MPC obtained from individual simulations were kept in close proximity to each other in a cubic box filled with water. At the start, the melittin dimer was moving far away from the MPC showing no interaction as shown in Fig. S2a. After some duration of simulation, MPC is near the N-terminal of melittin and the residues start to interact with the MPCs ligands as shown in Fig. S2b. After some time, Arg22 residue of MEL started to interact with the MPCs sulfur and gold atoms shown in Figs. S2c and S2d.

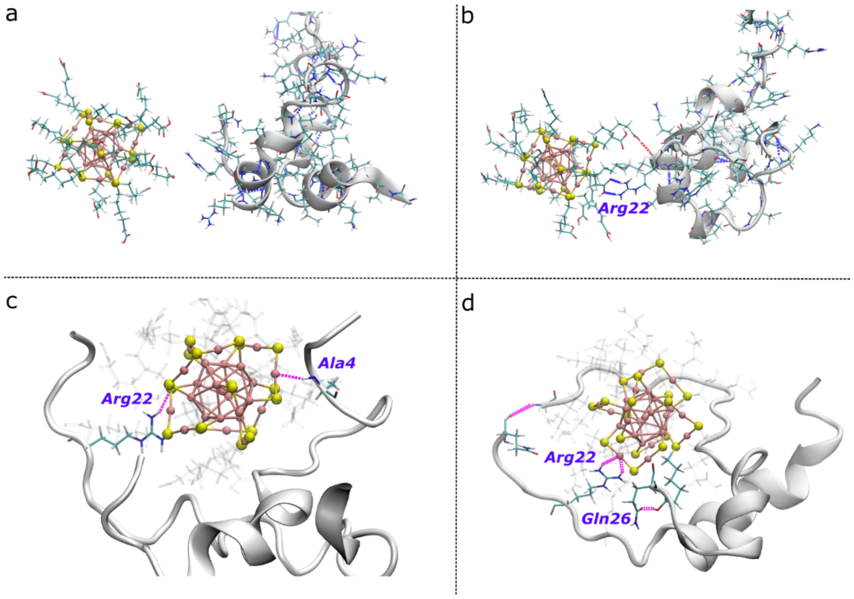


***Figure S2:*** *Interaction of Mel residues with MPC in dimer Mel:MPC(1:1) complex.*

From the RMSD analysis, we observed that MEL1 is fluctuating more and MEL2 is fluctuating less. After about 300ns of duration both the Melittin shows a constant interaction with minimal fluctuation. (Figure S3a) Through the RMSF analysis, we can see that the middle region of melittin residues shows minimum fluctuations this can be because of the hydrophobic patching of the hydrophobic groups of melittin residues such as Ile, Val, Ser, Leu, and Ala with the hydrophobic part of the ligands. (Figure S3b) Also, the N-terminal of one MELs fluctuates more while the other exhibit interactions with the MPCs ligands for some duration. The hydrogen bond analysis showed zero hydrogen bonds at the start of the simulation. (Figure S3d and S3e) This again can be the interpretation that MPC and dimer did not interact at the start of the simulation.

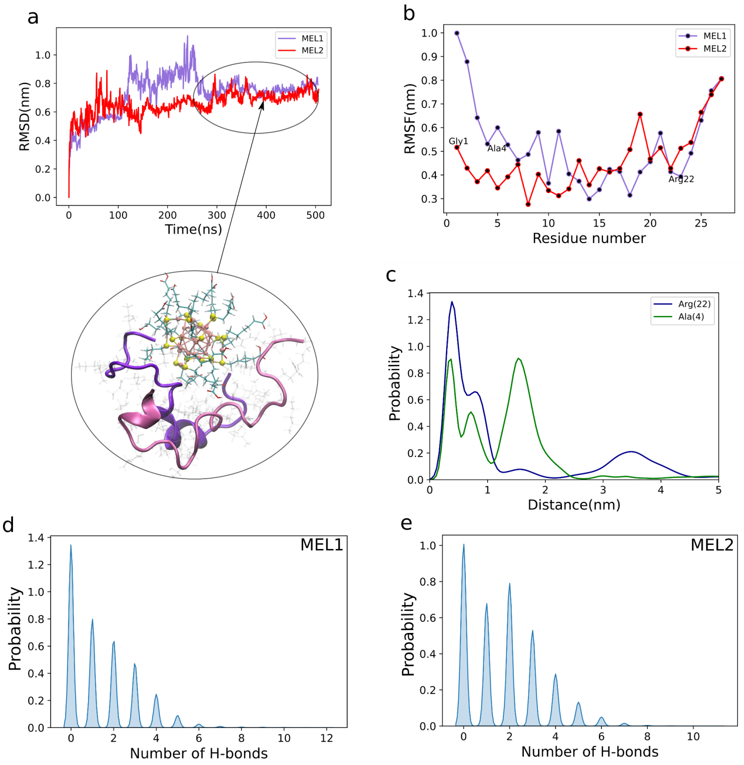


***Figure S3:*** *Simulation trajectory analysis of 1:1 Dimeric-Mel:$Au_{25}$-MPC system. a) RMSD analysis with a snapshot showing the stable region at the end of the simulation, b) RMSF analysis showing fluctuations in amino acid residues of Mel from their mean position during 500ns of simulation, c) Distance analysis of Arg22 and Ala4 interacting with MPC surface, d) Hydrogen bond analysis of Mel1 with MPC, e) Hydrogen bond analysis of Mel2 with MPC.*
